# Ginseng Ameliorates Effect of Prolonged Use of Omeprazole in Rat Renal Cortex through Decreasing Inflammation, Fibrosis and Apoptosis

**DOI:** 10.1101/2020.01.22.915587

**Authors:** Dina Ali Maher Abdel Dayem, Ahmed Sayed Mahmoud, Azza Hussein Ali, Nashwa Fathy Gamal El-Tahawy

## Abstract

Omeprazole is used in acid-related gastrointestinal disorders but has prolonged usage adverse effects. The aim was to study changes in renal cortex following chronic Omeprazole administration and the possible protective role of ginseng. Rats were divided into control (C-), Ginseng (G-), omeprazole (OM-), and omeprazole-ginseng (OM-G) groups. Serum urea and creatinine levels and 24-hours urine-protein were determined. Kidneys were processed for histological study. Serum urea and creatinine and 24-hours protein were significantly higher in OM-group compared to controls and significantly decreased in OM-G group comparing to OM-group. OM-group showed significant glomerular and tubulointerstitial injury with vascular congestion, inflammatory cell infiltration, partial or complete damage of apical brush border of most tubules, interrupted basement membranes of glomerular capillaries and tubules, marked increase in collagen deposition, and significant increases in COX-2 and caspase-3 immune-expression. Co-administration of ginseng with omeprazole resulted in marked and significant improvement of these morphological changes.

**Conclusion:** Omeprazole induced renal functional and morphological changes through inflammatory reaction, induction of fibrosis, cellular degeneration and apoptosis. Co-administration of ginseng ameliorated these effects through its anti-inflammatory, anti-fibrotic, and anti-apoptotic effects.

## 1. Introduction

The proton pump inhibitors (PPIs) are group of drugs that widely used to suppress gastric acid production by inhibiting functions of the H+/K+ ATPase pump in gastric mucosal parietal cells **(Al-Qaisi et al., 2018)**. Omeprazole, the prodrug of this family, is considered the first choice for most, if not all, acid related gastrointestinal disorders **(Nehra et al., 2018)**. It thought to has a good safety profile with a little frequency of adverse effects as nausea, diarrhea, headache and abdominal pain **(Al-Qaisi et al., 2018)** but it have been increasingly suspected of causing renal disorders particularly among older patients **(Fatima et al., 2019)** or even adolescents **(Noone et al., 2014)**. Several studies reported the hazardous effect of omeprazole on human kidney’s structure and function in meta-analytical studies **(Nochaiwong et al., 2017, Wu et al., 2018)**, retrospective case review **(Geevasinga et al., 2006)** and case reports **(Delve et al., 2003, Wall et al., 2000)**. However few studies have been done about omeprazole effect on the kidney of experimental animals in order to study the renal structural changes **(Mubeen et al., 2016)**.

Recently, the complementary and alternative medicines (CAM) have been rapidly growing. Ginseng, one of the different kinds of CAM is the most commonly used product **(Barnes et al., 2008)**. Ginseng contains numerous physiologically important elements, which included ginsenosides, fatty acids, peptides, alkaloids, polysaccharides, and other compounds **(Lee et al., 2010)**. Ginseng is potent antioxidant commonly used in herbal medicine. It is effective for reducing lipid peroxidation and subsequent tissue damage **(Chen and Huang, 2019, Mahmoud et al., 2014)**, and exhibited an anti-inflammatory properties **(Lee et al., 2015)**, anti-apoptotic and immunostimulatory effects **(Takeda and Okumura, 2018)**. The active metabolite of ginseng; ginsenoside, is reported to be effective in treatment of various disorders **(Shin et al., 2014)**. It has been found to protect from apoptosis and DNA fragmentation induced by chemicals and chemotherapeutic drugs encouraging protective efficacy **(Zhou et al., 2018)**.

Therefore, this study aimed to experimentally study the potential structural changes in the renal cortex of albino rats following chronic omeprazole administration and the possibly protective role of ginseng. Also, to identify the different mechanisms involved in these changes.

## 2. Results

### 2.1. Biochemical results

Rats of the G-group had insignificant increases in serum urea, serum creatinine and the total 24-hours urinary proteins compared to the C-group (all p>0.05). The OM-group and OM-G group had significant increases in serum urea, serum creatinine and the total 24-hours urinary proteins compared to the C-group and G-group (all p<0.001). However, co-administration of ginseng in OM-G group significantly decreased these levels compared to OM-group (all p<0.001) (Table 1).

**Table 1:**
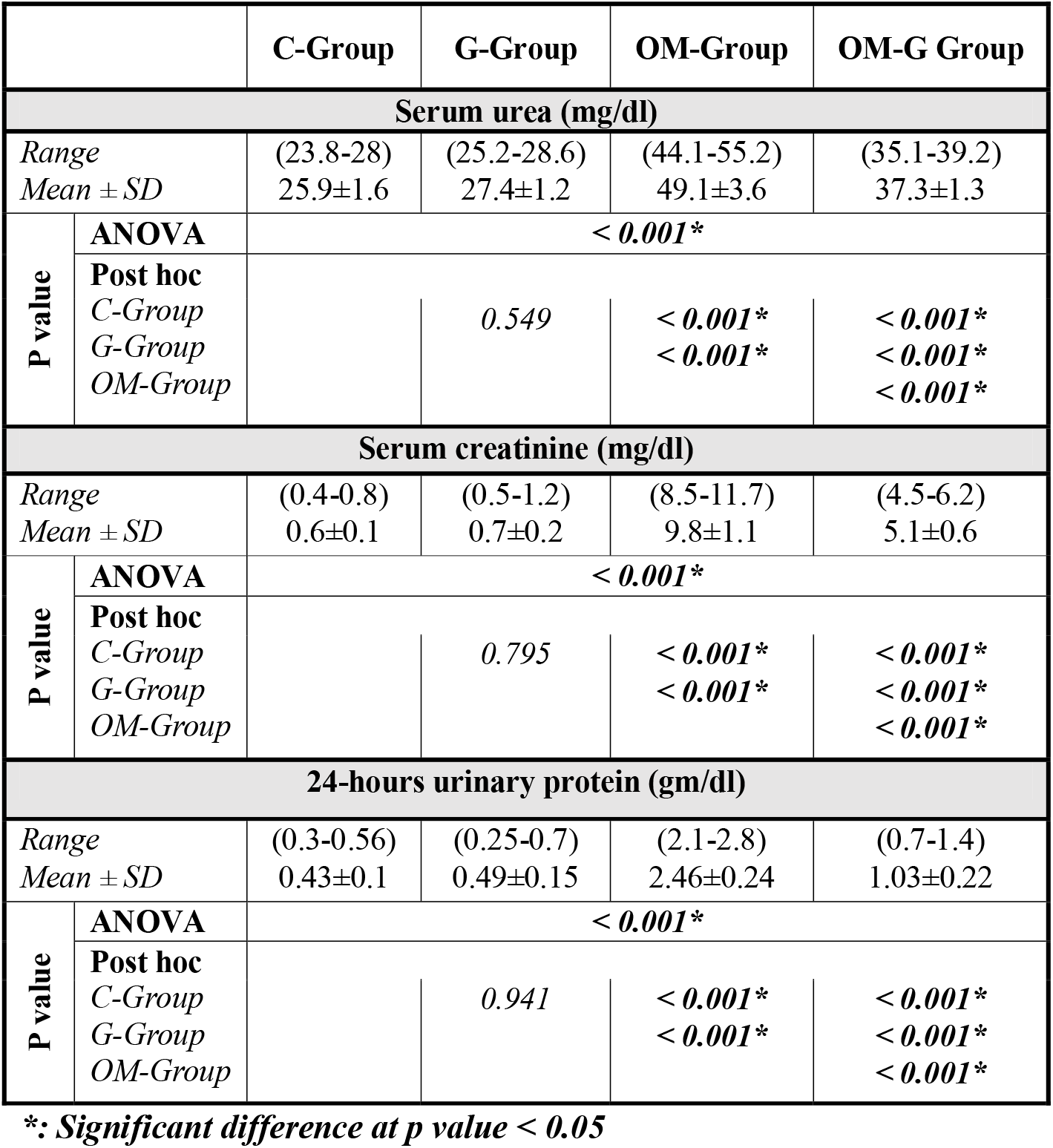
Serum urea (mg/dl), Serum creatinine (mg/dl) and 24-hours urinary protein (gm/dl) levels in the studied groups (n=8)

### 2.2. H&E results

The C-group and G-group had a normal histological structure of the renal cortex. The cortices were formed of renal corpuscles, proximal (PCTs) and distal (DCTs) convoluted tubules. The renal corpuscles were formed of the glomerular tuft of capillaries surrounded by Bowman’s capsule. The DCTs had larger regular distinct lumina than the PCT. Macula densa cells of the DCTs were in close proximity to the renal corpuscle (Fig. 1). The OM-group showed histological changes in the renal cortices demonstrating significant glomerular and tubulointerstitial injury with obvious vascular congestion. Congestion of peritubular capillaries and interstitial hemorrhage were observed. Renal corpuscles appeared congested, distorted or even shrunken and atrophied. Also, there was marked glomerular vacuolation with widening of the Bowman’s space. Marked tubular distortion was observed; most cells showed vacuolar degeneration, some cells had ghosts of nuclei with disappearance of cytoplasm and others showed apoptotic figures (deeply acidophilic cytoplasm and dark pyknotic nuclei). Epithelial flattening with reduction of cell height and desquamation of some cells with cellular debris in the lumina were also evidenced. There was interstitial mononuclear inflammatory cellular infiltration composed mostly of lymphocytes. While administration of omeprazole and ginseng in combined form in the OM-G group ameliorated the damaging effects of omeprazole as clearly noticed by the reduction of histological changes compared to the OM-group. The renal corpuscles appeared more or less normal with mild congestion of glomerular capillaries and decreased peritubular congestion with absence of interstitial hemorrhage. Minimal intraluminal hyaline casts were observed with no cellular casts in most tubules. Most PCTs and DCTs retained their apparently normal lining epithelium; simple cubical epithelium (Fig.2). This was confirmed morphometrically in Table 2.

**Table 2:** Histopathological Scoring of morphological changes in the studied groups (H& E stained sections) (n=8)

| **Findings** | **C- Group** | **G- Group** | **OM- Group** | **OM-G Group** |
| --- | --- | --- | --- | --- |
| Interstitial: |  |  |  |  |
| -Hemorrhage | - | - | +++ | + |
| -Inflammatory cells | - | - | +++ | - |
| Renal corpuscles: |  |  |  |  |
| -Glomerular vaculation | - | - | ++ | - |
| -Shrunk / distorted renal corpuscles | - | - | ++ | - |
| Renal Tubules: |  |  |  |  |
| -Tubular cells vaculation | - | - | +++ | +/- |
| -Lumen widening and Distortion | - | - | +++ | +/- |
| -Casts | - | - | ++ | + |

Normal (-), in-between normal and mild (+), mild (++), moderate (+++) and sever (++++).

**Fig. 1:**
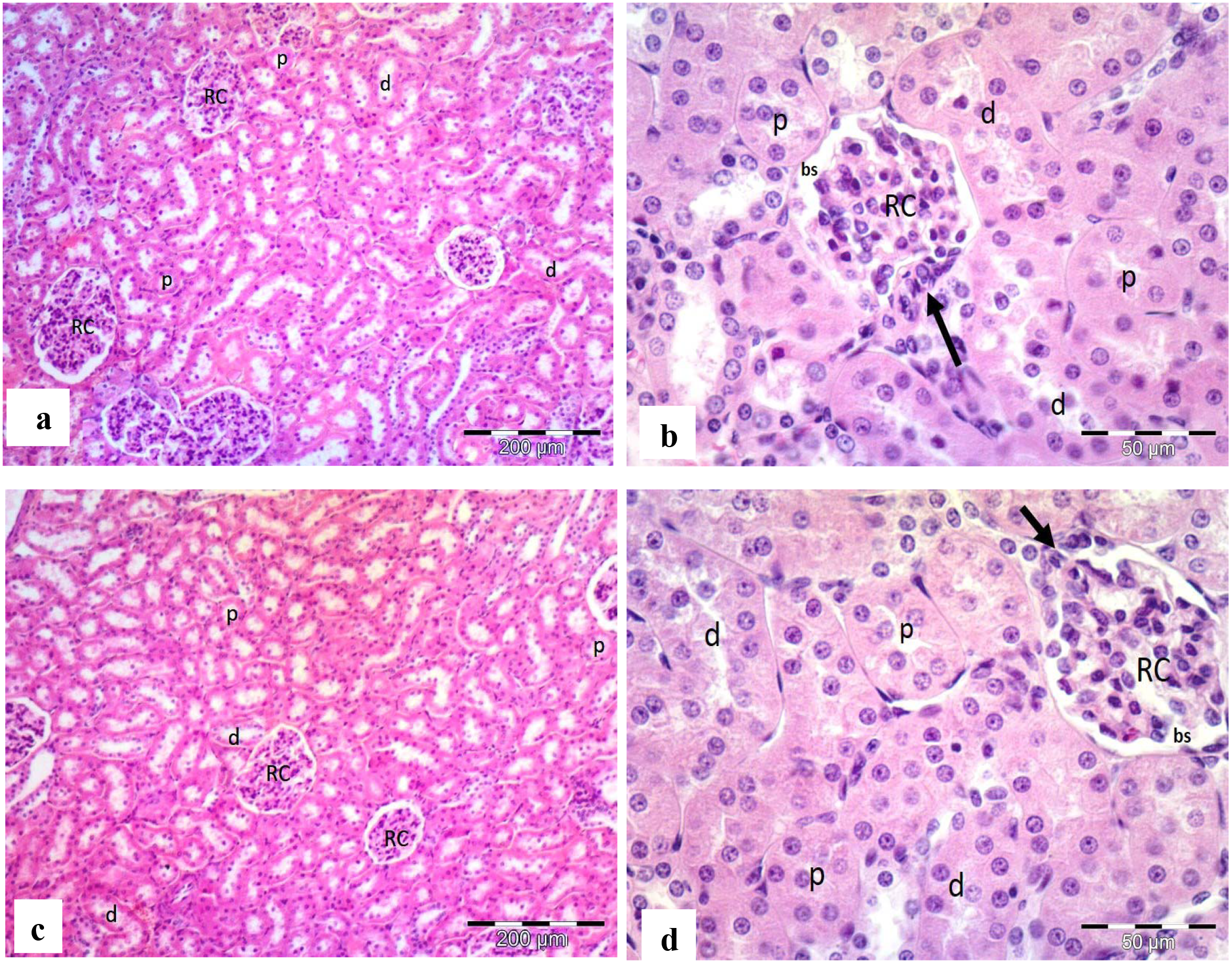
Photomicrographs of renal cortex from **C-group** (a,b) **and G-group** (c,d) showing normal lobular organization of the renal cortex consisting of renal corpuscles (RC), PCTs (p), and DCTs (d). Notice the Bowman’s space (bs), macula densa cells (arrow) of DCT and the regular distinct lumena of convoluted tubules. H&E aX100; bX400

**Fig. 2:**
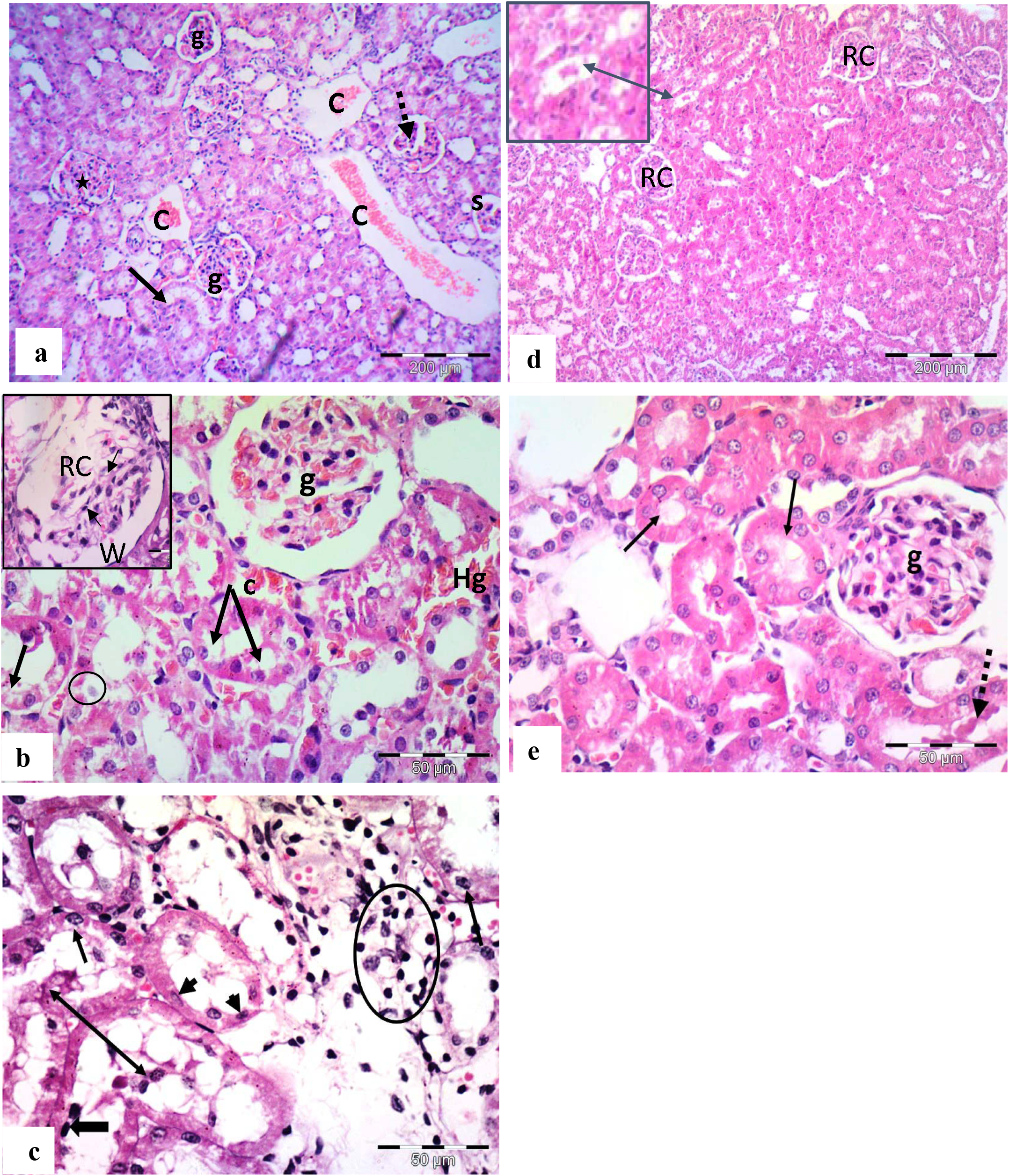
photomicrographs of renal cortices of **OM-group** (a-c) **and OM-G group** (d,e) showing: (a) Congested glomerular capillaries (g), interstitial vascular congestion (C) and vacuolation of tubular cells (arrow). Notice the shrunken (s), distorted (dotted arrow) and hypertrophied renal corpuscle (star). (b) Congestion of peritubular capillaries (c) and interstitial hemorrhage (Hg). Notice vacuolation of some tubular cells (arrows) and other cells with ghosts of the nuclei and disappearance of cytoplasm (circle). Inset shows deformed renal corpuscle (RC) with widening of Bowman‘s space (W). (c) Tubular cell distortion with flattening (thin arrows), desquamation with cellular debris in lumina (double head arrow), or apoptotic figures (thick arrow). Notice inflammatory cell infiltrate mostly lymphocytes (circle). (d) & e) Restoration of apparently normal cortex of the **OM-G group**, only minimal intraluminal hyaline casts (double head arrow) and mild congestion of glomerular capillaries (g).Most PCTs and DCTs retained their apparently normal simple cubical epithelium (arrows). H&E a,dX 100; b, c and eX 400

### 2.3. The PAS staining results

Renal cortical tissues of C-group and G-group showed normal positive PAS reactions in the basement membranes of intact glomerular blood capillaries, parietal layers of Bowman’s capsules, PCT and DCT cells. It was also observed in the intact apical brush borders of PCT and DCT cells. The OM-group showed reduced PAS reaction in the form of partial or complete loss of the brush border of most distorted tubules. The basement membranes of renal tubules and glomerular capillaries were also seen interrupted and appeared thin at some sites. The OM-G group showed preserved reaction of most brush borders of PCTs and DCT and basement membranes which appeared continuous and intact. Only few renal tubules were seen in some sections with reduced PAS reaction in their brush borders (Fig.3, Table 3).

**Table 3:** Scoring of Fibrotic changes in the studied groups (Masson trichrome stained sections) (n=8)

| Findings | C- Group | G- Group | OM- Group | OM-G Group |
| --- | --- | --- | --- | --- |
| Interstitial fibrosis stain | 0 | 0 | 3 | 1 |

Score 0 : <10%, Score 1: 10–25%, Score 2: 26%–50% and Score 3: >50% of renal cortex involved by interstitial fibrosis.

**Fig. 3:**
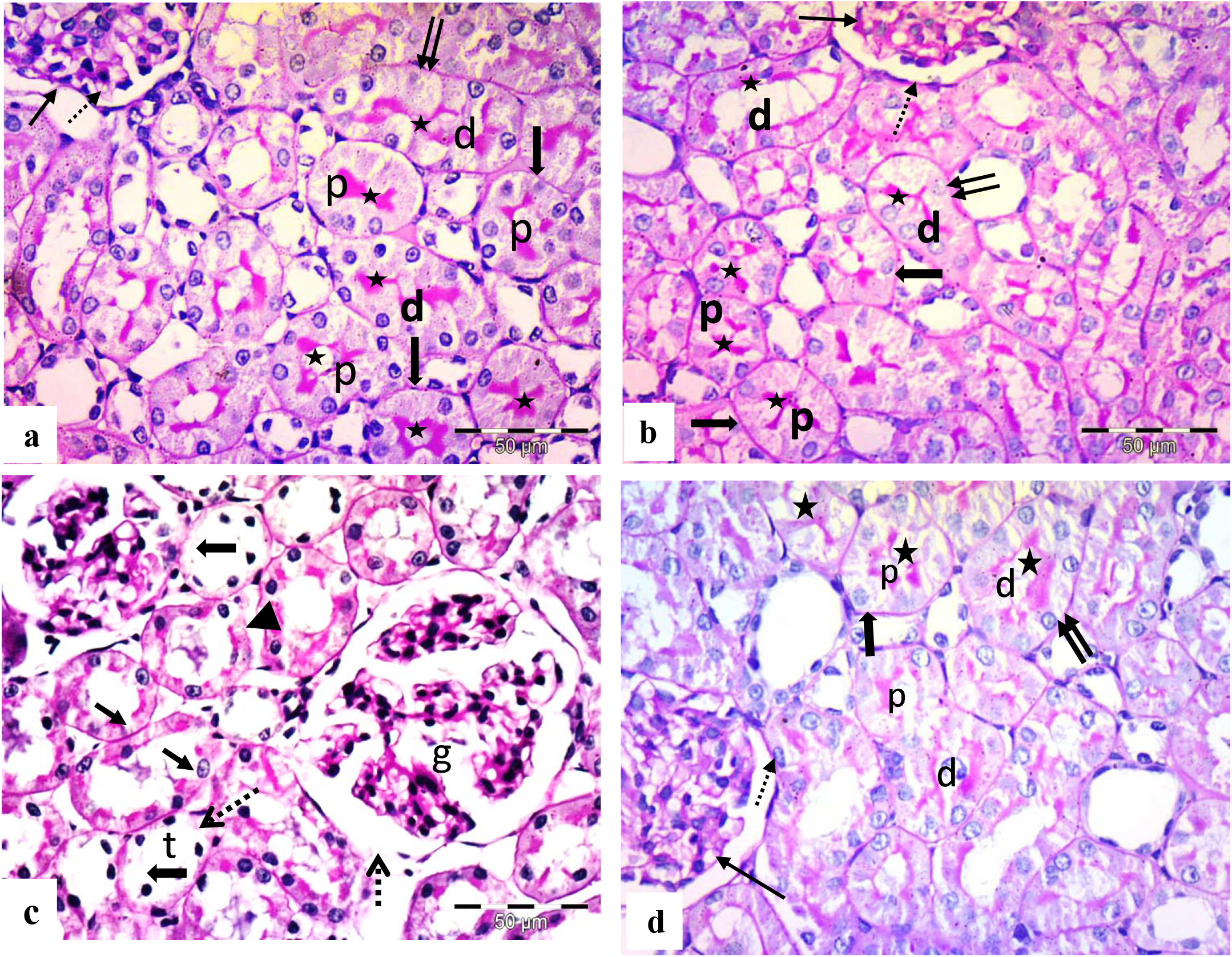
Photomicrographs of renal cortices from: (a) **C-group** showing positive PAS reaction in the BMs of glomerular blood capillaries (thin arrow), parietal layers of Bowman’s capsules (dotted arrow), PCTs (thick arrows) and DCTs (double arrows) cells. Notice the staining of brush borders (stars) of the PCT (p) and DCT (d). (b) **G-group** showing the same histological finding as **C-group**. (c) **OM-group** showing partial (thin arrows) or complete loss (thick arrows) of the brush border of distorted renal tubules. Notice interruption of basement membrane (dashed arrow) of the distorted glomerulus (g) and tubules (t). (d) **OM-G group** showing preserved reaction of most brush borders (stars) of PCTs (p), DCTs (d), continuous basement membranes of glomerular blood capillaries (thin arrow), parietal layers of Bowman’s capsule (dotted arrow), PCTs (thick arrows) and DCTs (double arrows) cells. PAS X400

### 2.4. Masson trichrome stain results

The C-group and G-group showed scanty fine strands of collagen fibers around the renal calyces, surrounding the glomeruli and among the renal tubules. The OM-group showed increased amount of fibrosis around: the renal calyces, the glomeruli and among the tubules compared to the other groups. While, the OM-G group showed reduction of the amount of collagen fiber deposition around the renal calyces, surrounding the glomeruli and among the tubules compared to OM-group to become nearly similar to normal (Fig.4, Table 4).

**Table 4:**
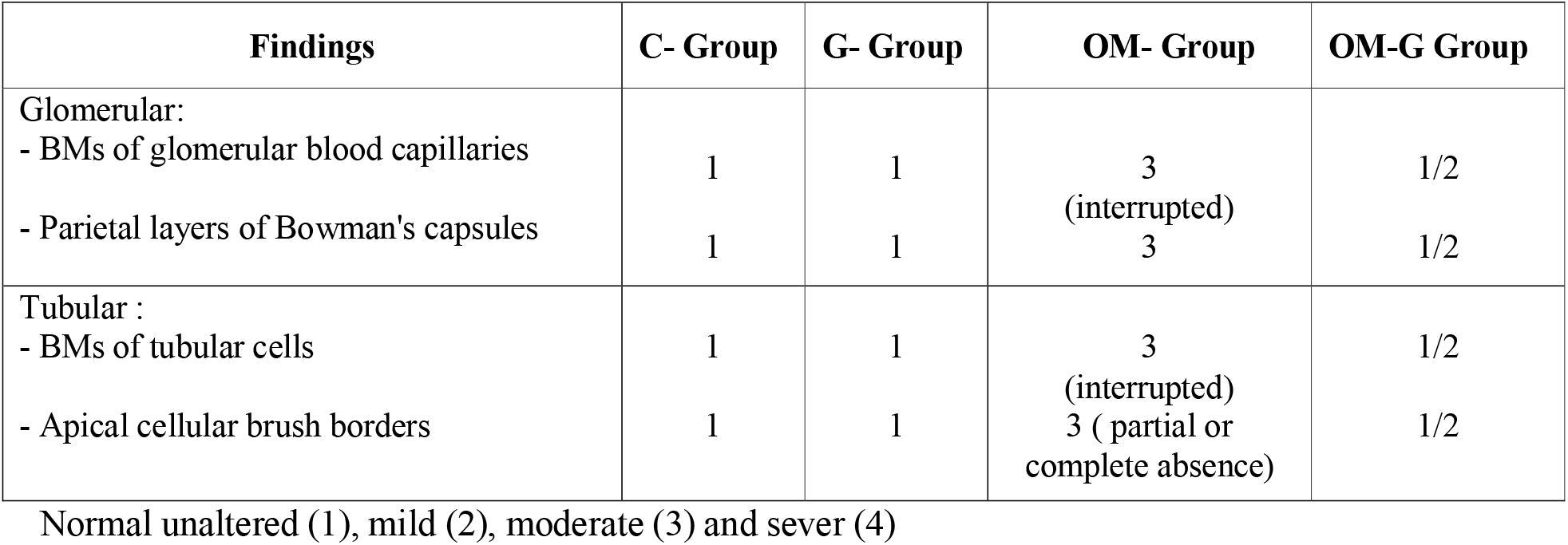
Scoring of morphological changes in the studied groups (PAS stained sections) (n=8)

| Findings | C- Group | G- Group | OM- Group | OM-G Group |
| --- | --- | --- | --- | --- |
| Glomerular: |  |  |  |  |
| - BMs of glomerular blood capillaries | 1 | 1 | 3<br>(interrupted) | 1/2 |
| - Parietal layers of Bowman's capsules | 1 | 1 | 3 | 1/2 |
| Tubular : |  |  |  |  |
| - BMs of tubular cells | 1 | 1 | 3<br>(interrupted) | 1/2 |
| - Apical cellular brush borders | 1 | 1 | 3 (partial or complete absence) | 1/2 |

Normal unaltered (1), mild (2), moderate (3) and sever (4)

**Fig. 4:**
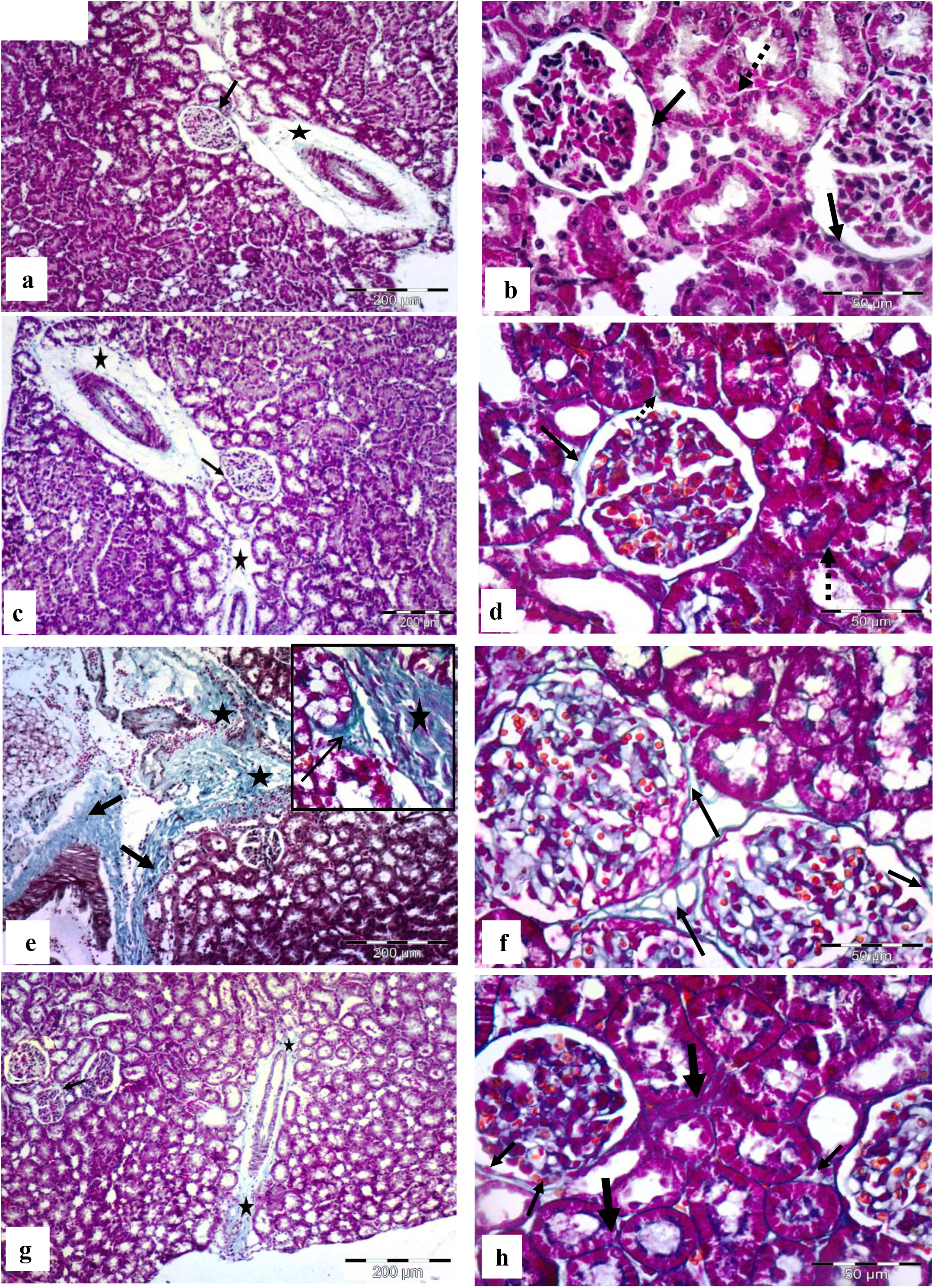
Photomicrographs of renal cortices from: (a&b) **C-group** showing scanty fine strands of collagen fibers surrounded the glomeruli (arrow), around the renal calyx (star) **(b)** and between the renal tubules (dotted arrow). (c&d**) G-group** showing the same collagen fibers distribution as in C-group. (e&f) **OM-group** showing increased amount of collagen fibers around the renal calyx (thick arrows) and in between lobules (stars). **(f)** increased amount of collagen fibers surrounding the glomeruli (arrows). Increased amount of collagen fibers between lobules (star) and creeping between tubules (arrows). (g&h) **OM-G group** showing reduction of collagen fibers: surrounding the renal calyx (stars) and the glomeruli (arrow)**. (h)** and between tubules (thick arrow) compared to **OM-group**. Masson trichrome a,c.e,g X100; b,d,f.h X400

### 2.5. Immunohistochemical results

#### Regarding COX-2 immune-staining

The C-group and G-group showed insignificant difference in the area fraction of COX-2 expression (p=0.998) which appeared as faint cytoplasmic expression in few macula densa cells. The OM-group showed significant increase in the area fraction of COX-2 compared to the other 3 groups (all p<0.001). It displayed numerous deeply stained COX-2 expressions in macula densa, PCT, DCT and some glomerular cells. However, there was a significant decrease in the OM-G group compared to OM-group (p<0.001) and the expression seen only in macula densa cells (Fig.5, Table 5).

**Table 5:** Mean area fraction (%) of COX-2 and Caspase-3 in the studied groups (n=8)

|  | C-Group | G-Group | OM-Group | OM-G Group |
| --- | --- | --- | --- | --- |
| **COX -2 surface area fraction (%)** |  |  |  |  |
| Range | (17.8-20.7) | (17.4-20.3) | (40.1-43) | (25.5-30.8) |
| Mean ± SD | 19.2±1 | 19.1±1 | 41.7±1.1 | 28.5±1.9 |
| **P value** | **ANOVA** | **< 0.001\*** |  |  |
|  | **Post hoc** |  |  |  |
|  | C-Group | 0.998 | < 0.001* | < 0.001* |
|  | G-Group |  | < 0.001* | < 0.001* |
|  | OM-Group |  |  | < 0.001* |
| **Caspase-3 surface area fraction (%)** |  |  |  |  |
| Range | (0.1-1.2) | (0.7-1.3) | (24.4-28.5) | (10.1-15.8) |
| Mean ± SD | 0.9±0.3 | 1±0.2 | 26.2±1.8 | 13.4±2.2 |
| **P value** | **ANOVA** | **< 0.001\*** |  |  |
|  | **Post hoc** |  |  |  |
|  | C-Group | 0.996 | < 0.001* | < 0.001* |
|  | G-Group |  | < 0.001* | < 0.001* |
|  | OM-Group |  |  | < 0.001* |

\*: Significant difference at p value < 0.05

**Fig. 5:**
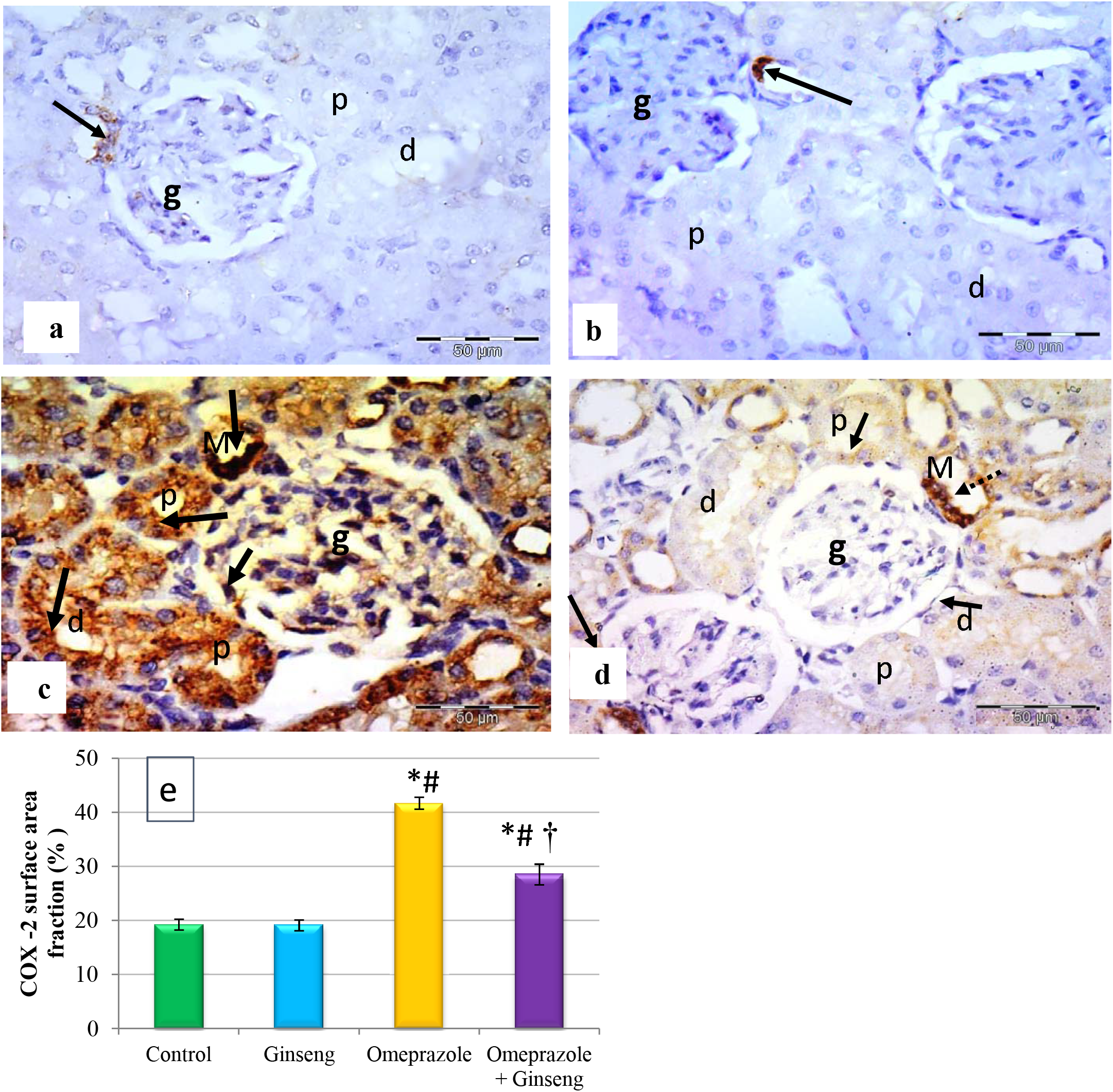
Photomicrographs of immune-stained renal cortices for COX-II from: (a) **C-group** showing COX-2 few immune-expression only in macula densa cells (arrow). Notice the negative immune-reactivity in the glomerular (g), PCT (p) and DCT (d) cells. (b) **G-group** showing few immune-expression only in macula densa cells (arrow). (c) **OM-group** showing increased COX-2 immune-expression (arrows) in the PCT (p), DCT (d) cells, macula densa cells (M) of DCT and some glomerular cells (g). (d) **OM-G group** showing decreased COX-2 immune-expression (arrows) in most PCT (p) and DCT (d) with deep expression in macula densa cells (M) (dotted arrow). Immunohistochemistry, counterstained with H: X400 (e) The mean area fraction of COX-2 immune-reactivity in the studied groups (n=8). Significant: * vs C-group, # vs **G-group**, and † vs **OM-group**.

#### Regarding activated caspase-3 immune-staining

The C-group and G-group showed no detectable immunolabeling for activated caspase-3 with insignificant difference in the area fraction (p=0.996). The OM-group showed high immune-reactivity in renal corpuscular and tubular cells with a significant increase compared to the other 3 groups (all p<0.001). The OM-G group showed significant decreased immune-reactivity compared to the OM-group (p<0.001) (Fig.6, Table 5).

**Fig. 6:**
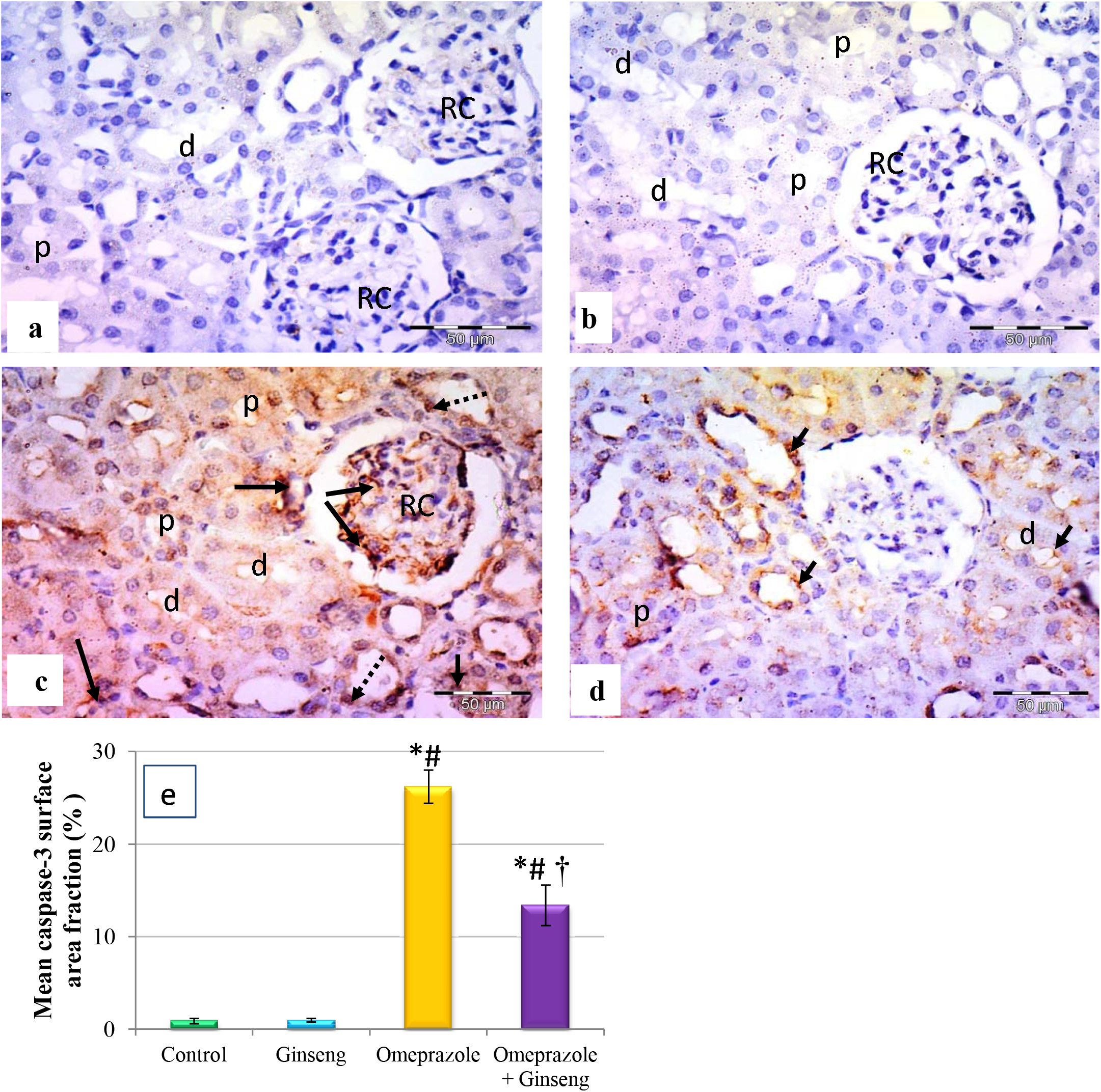
Photomicrographs immune-stained renal cortices for caspase-3 from: (a) **C-group** showing negative caspase-3 immune reactivity in the cells of the renal corpuscle (RC), PCT (p) and DCT (d). (b) **G-group** showing negative immune-reactivity as **C-group**. (c) **OM-group** showing positive cytoplasmic immune-expression (arrows) in the cells of the renal corpuscle (RC), PCT (p) and DCT (d). Notice some nuclear immune-expression (dotted arrows). (d) **OM-G group** showing decreased caspase-3 expression with faint expression in some tubular cells (arrows). Immunohistochemistry, counterstained with H: x400 **e)** The mean area fraction of COX-2 immune-reactivity in the studied groups (n = 8). Significant: * vs **C-group**, # vs **G-group**, and † vs **OM-group**.

## 3. Discussion

In this study, serum urea and creatinine levels showed marked increases after omeprazole administration which was in line with the observations of previous human case studies who found the same in their patients **(Noone et al., 2014, Wall et al., 2000)**. Interstitial inflammation can induce accumulation of extracellular matrix leading to impairment of renal functions **(Rossert, 2001)**. Also, the 24-hours urinary protein level had a significant increase in OM-group, this was in agreement with other authors **(Berney◻Meyer et al., 2014**, **Geevasinga et al., 2006)**. Inflammatory mediators causes endothelial cell activation which lead to vascular permeability and subsequently proteinuria **(Niklasson et al., 2010)**. So the inflammation and structural changes observed in OM-group in this study, might attributed to the impaired renal functions. However, co-administration of ginseng and omeprazole in OM-G group resulted in significant improvement in kidney functions evidenced by reduction in serum urea, serum creatinine and the 24-hours urinary protein levels.

Omeprazole administration caused marked variable and patchy morphological changes demonstrating significant glomerular and tubulo-interstitial injury which was in accordance with results of other researchers **(Mubeen et al., 2016)**. They stated that omeprazole has toxic effects in the vasculature of the kidney which evidenced by glomerular congestion and atrophy in addition to the stromal hemorrhage and congestion resulting from weakness of renal vasculature structure. In addition, accumulation of inflammatory cells and the generation of cytokines from damaged tubules resulted in glomerulo-tubular disconnection and collapse of glomeruli **(Chevalier and Forbes, 2008)**. However, these results disagreed with the results of other studies **(Brewster and Perazella, 2007a, Geevasinga et al., 2006, Sampathkumar et al., 2013)** who reported that almost uniform renal biopsy findings of extensive lymphocytic infiltrations involving the interstitium but with sparing of the glomeruli. In this study, inflammatory mononuclear cellular infiltration mainly of lymphocytes was evident which was in accordance with other studies **(Sethi et al., 2017, Muriithi et al., 2014, Noone et al., 2014)**, meanwhile others mentioned that the interstitial inflammation includes inflammatory cells of several types **(Silva, 2004)**.

The glomerular and tubular cell cytoplasmic vacuolization observed in this work could be attributed to cytoplasmic degeneration, necrosis and disturbed ion transport system **(Condron et al., 1994)** or occurs as a cellular defense mechanism against injurious materials which segregated in vacuoles interfering with cellular metabolism **(Robbins et al., 2011)**.

Omeprazole is metabolized by cytochrome P-450 so it has a high affinity to cortical tubular cells which predominantly contain cytochrome P-450 **(Brewster and Perazella, 2007b)** leading to the release of free oxygen radical (ROS). These ROS covalently bind to cellular macromolecules especially polyunsaturated fatty acids, initiating autocatalytic lipid peroxidation, DNA damage, protein degeneration and apoptosis **(Al-Olayan et al., 2014, Hesami et al., 2014)**. These might interpreted the tubular degenerations and the apoptotic figures observed in the tubular cells in the OM-group.

There is an interesting reciprocal relationship between proteinuria and tubular injury. The local infiltration of leukocytes together with activation of tubular and interstitial cells initiate the release of inflammatory cytokines and growth factors, e.g. the transforming growth factor-β (TGF-β), vascular endothelial growth factor-A (VEGF-A). This appears to be a major cause of nephrin down regulation and occurrence of proteinuria **(Tanaka and Nangaku, 2011)**. Nephrin, which considered an important protein of the slit diaphragm of podocyte, has anti-apoptotic signaling properties. In addition, TGF-β1 causes podocyte apoptosis and increase in extracellular matrix formation and deposition **(Wolf and Ziyadeh, 2007)**. On the other hand, there was experimental evidence that the filtered macromolecules provoke harmful effects on tubular cells as energy depletion and lysosomal rupture leading to a cascade of events that culminates in cell injury **(Kato et al., 2008, Nangaku et al., 2002, Tang et al., 2002)**. So, it appeared to be a vicious circle. Co-administration of ginseng with omeprazole resulted in amelioration of the damaging effects of omeprazole. These results might be due to its antioxidant property. It acts as a free-radical scavenger and lipid peroxidation inhibitor **(Zhang et al., 2010)**. This ameliorative effect of ginseng might be also due to its ability to bind to glucocorticoid receptor triggering an activation of glucocorticoid response which encourage cell proliferation and improves the survival rate of new formed cells **(Shen and Zhang, 2003)**.

The PAS reaction in OM-group was reduced with partial or complete damage of the brush border of most distorted tubules and interrupted basement membranes of kidney tubules and the glomerular capillaries. This was in accordance with results of the study of Berney◻Meyer and his colleagues **(Berney◻Meyer et al., 2014)**. Peroxidation of lipids is evident in the renal brush border and dramatically changes the properties of the biological membranes leading to severe cell injury with loss of its apical brushing **(Adewole et al., 2007, Al-Yahya et al., 2013)**. Ginseng markedly improve these changes which was in the same line with others who suggested that ginseng had a protective effect through protection of cells and tissues from the destructive effects of the free radicals **(Zidan RA, 2015)**. This effect could be also attributed to the previously mentioned lipid peroxidation inhibitory effect of ginseng **(Zhang et al., 2010)**.

In Masson trichrome stained sections, there was marked increase in collagen fibers deposition in OM-group which was in accordance with other researchers **(Sethi et al., 2017, Muriithi et al., 2014)**. These finding could be explained as inflammatory cells might be the physiopathological link between tubular epithelial cell injury and renal fibrosis. Experimental studies and clinical observations proved that the process of fibrosis occurs due to an altered crosstalk between the tubular cells and the interstitial fibroblasts **(Chawla et al., 2011)**. Macrophages and the inflammatory cells are important sources of cytokines and growth factors which in turn play a critical role in the progress of interstitial fibrosis **(Tanaka and Nangaku, 2011)**. A previous explanation for increased fibrous deposition discussed a phenomenon of “Epithelial - Mesenchymal Transition” observing that a considerable number of the interstitial fibroblasts, originated from the neighboring tubular cells. Hence and under certain pathological conditions, active interstitial fibroblasts can be generated from tubular epithelial cells leading to more production of the extracellular matrix **(Iwano et al., 2002)**. Coadministration of ginseng and omeprazole in the OM-G group resulted in reduction of the amount of collagen fiber deposition to become nearly similar to normal. This finding was in the same line with other research results which stated that Panax ginseng is effective against CCl4-induced fibrogensis through prevention of CCl4-oxidative damage in liver **(Karadeniz et al., 2009)**.

Cyclooxygenase-2 (COX-2), an enzyme responsible for formation of important biological mediators including prostaglandins, prostacycline and thromboxane; is undetectable in most normal tissues. However, it could be an inducible enzyme in most tissues exposed to inflammation **(Warner and Mitchell, 2002)**. In response to an insult tubular cells can become activated and produce pro-inflammatory molecules such as cytokines, growth factors, adhesion molecules, or chemokines resulting in inflammatory response **(Rossert, 2001)**. In the current study, immune-histochemical detection of COX-2 restricted to macula densa cells in the control group. This was explained as COX-2 in rat Macula densa cells was related to renin release **(Kömhoff et al., 2000)**. On the other hand, there was significantly increase in COX-2 expression in OM-group which not restricted to macula densa cells but also extend to display deeply stained cytoplasmic expression in PCT, DCT and some glomerular cells due to the observed lymphocytic infiltration. Omeprazole induced renal injury as an idiosyncratic immune reaction secondary to a cell-mediated process in the kidney **(Krishnan and Perazella, 2015)**. In the current study, the decreased inflammatory cell infiltration and the significant decrease of COX-2 immune-expression in the OM-G group indicated anti-inflammatory effects of ginseng.

Evidences of apoptotic process were clearly observed in OM-group and ginseng treatment decreased it significantly. Omeprazole found to decrease levels of the anti-apoptotic proteins (Bcl-2 and Bcl-XL) **(Patlolla et al., 2012)** which control cell survival through inhibition of the intrinsic pathway of apoptotic including caspase-3 cleavage **(Youle and Strasser, 2008)**. Tubular epithelial cell apoptosis contributes to tubulointerstitial fibrosis **(Zhou et al., 2014)**. Ginseng has a potent anti-apoptotic effect as it increases the expression of the anti-apoptotic gene; Bcl-2 which antagonize the previously mentioned mechanism of omeprazole induced apoptosis **(Kim et al., 2010)**. Also, ginseng saponin decrease cisplatin induced cytosolic free Ca^2+^ ions overload and formation of DNA-protein cross-link **(Abdel-Wahhab and Ahmed, 2004)**. These could explain the significant decrease of caspase-3 expression in this study.

In conclusion, omeprazole induced renal functional and morphological changes through immune mediated inflammatory reaction, induction of fibrosis, cellular degeneration and apoptosis. Co-administration of ginseng ameliorated these effects through its antiinflammatory, antifibrotic, and antiapoptotic effects.

## 4. Material and method

### 4.1. Ethics

The experiment underwent approval by the ethical committee for animal handling for research, Faculty of Medicine, Minia University according to international guidelines (Act 1986).

### 4.2. Chemicals

- Omeprazole (Pepzol MR®) tablets containing 20 mg of omeprazole were obtained from Hekma Pharmaceuticals (Cairo, Egypt). Tablets were freshly dissolved in 100 ml distilled water and was used within 5 minutes of preparation with the dose of 20 mg/kg by oral gavage.
- Ginseng (GINSANA®) capsules contained dried roots of Panax ginseng. Each capsule had 100 mg of ginseng. Capsules were obtained from EPICO Pharmaceuticals (Cairo, Egypt). They were freshly dissolved in 100 ml distilled water and was used within 5 minutes of preparation with the dose of 100 mg/kg by oral gavage.
- COX2 antibody (monoclonal rabbit antibody), ultra-vision one-detection system, HRP Polymer and diaminobenzidine tetra hydrochloride (DAB) plus Chromogen (Thermo Fisher Scientific, USA). Code NO. RM-9121-R7 purchased from Sigma Aldrich Company, Egypt.
- Anti-cleaved caspase-3 antibody (polyclonal rabbit antibody), ultra-vision one detection system, HRP Polymer and DAB+Chromogen (Thermo Fisher Scientific, USA). Code NO. PA1-29157 purchased from Sigma Aldrich Company, Egypt.

### 4.3. Animals

Thirty two male albino rats (8-12 weeks, 150-200 gram) were purchased from the Faculty of Agriculturés animal house, Minia University. Rats were housed in clean open to the room environment plastic cages with normal light/dark cycles. All animals became acclimatized for 2 weeks before the beginning of the study and fed standard laboratory diet and water ad libitum.

### 4.4. Experimental design

Rats were equally and randomly divided into 4 groups (8 rats for each group):

1. **Control Group (C-group):** received standard rat chow diet and clean water.
2. **Ginseng Group (G-group):** received 100 mg/kg of ginseng daily in a single dose dissolved in 100 ml distilled water by oral gavage for 12 weeks **(Zidan RA, 2015)**.
3. **Omeprazole Group (OM-group):** received 20 mg/kg omeprazole daily in a single dose dissolved in distilled water by an oral gavage for 12 weeks **(Shah et al., 2016)**.
4. **Omeprazole-Ginseng Group (OM-G group):** received a daily dose of ginseng (100 mg/kg/day) and omeprazole (20 mg/kg body weight/day) both dissolved in distilled water and given simultaneously by an oral gavage for 12 weeks.

### 4.5. Biochemical study

- **Determination of 24-hours protein in urine:** Rats were separated singly in metabolic cages for 24-hours urine collection before the end of the experiment and urine proteins were measured **(Abuelo, 1983)**.
- **Determination of serum urea and creatinine levels:** At the end of the experiment, blood samples were drawn through cardiac puncture with needle mounted on 5-ml syringe technique. The blood samples were allowed to stand for about 30 minutes to clot and centrifuged for 15 min. Sera were separated and used to determine the levels using ready to use creatinine **(Fossati et al., 1983)** and urea kits **(Tietz, 1995)**.

### 4.6. Histological procedures

Rats from all groups were decapitated after 12 weeks under light halothane anesthesia. Kidneys were rapidly removed, rapidly fixed in 10% formalin, paraffin-embedded, sectioned (5-μm thick) then deparaffinized, hydrated and washed for:

- Staining with hematoxylin & eosin (H&E), Periodic acid Schiff’s reaction (PAS), and Masson trichrome stains according to Suvarna et al., **(Suvarna et al., 2018)**.
- Immunohistochemistry according to de Alcântara et al., **(de Alcântara et al., 2017)**: in brief, the endogenous peroxidases were quenched by H_2_O_2_ and washed in tris buffer saline. Sections were incubated with the diluted 1^ry^ antibodies for cleaved caspase-3 (1:200) and the ready used COX-2 antibody. Subsequently it was incubated in biotinylated goat anti-rabbit 2ry antibodies (1:1000). Then incubated with Vectastain ABC kits (Avidin, Biotinylated horse radish peroxidase Complex), DAB was added for 5-10 min. The immune-positivity appeared as brown cytoplasmic reaction for Cox-2, and cytoplasmic, nuclear or both for caspase-3.

Tissues of colonic carcinoma used as the positive control for COX-2 and tonsils were used as positive control for activated caspase-3. The negative control sections were obtained by removal of the 1^ry^ antibody (figures not included).

### 4.7. Image capture

Olympus light microscope and its Olympus digital camera (Olympus, Japan) were used for examination of sections and capturing images. Images were saved as jpg.

### 4.8. Morphometrical study

- **Histopathological scoring:** The frequency and severity of histopathological lesions in the kidney were assessed semi-quantitatively in 10 non overlapping field in H&E sections **(Saber et al., 2013)**: Score - : assigned normal, Score +: in between normal and mild level, Score ++: (mild level) less than 25 %, Score +++: (moderate level) less than 50 %, and Score ++++: (severe level) less than 75 % of the total fields examined revealed histopathological alterations.
- **Masson trichrome findings for scoring of fibrosis (Sethi et al., 2017):** Score 0 : <10%, Score 1: 10–25%, Score 2: 26%–50%, and Score 3: >50% of renal cortex involved by interstitial fibrosis.
- **PAS findings**: alterations of parietal cells of Bowman’s capsule, basal membrane, and tubular cells **(de Souza et al., 2009):** Score 1: Unaltered characteristics, Score 2: Mild alterations of parietal cells, minimal interruption basal membrane and mild interruption of brush border and basal membrane, Score 3: Moderate alterations of parietal cells, moderate interruption basal membrane and moderate interruption of brush border and basal membrane, and Score 4: Marked alterations of parietal cells, intense interruption of basal membrane and complete interruption of brush border and basal membrane.
- **Measuring area fraction of COX-2 and caspase-3 immune-reactivity:** Image J 22 software (open source Java image processing program) was used for measuring the area fraction of COX-2 and caspase-3 immune-reactivity. Area fraction was measured in a standard measuring frame/5 photomicrographs for each group X 400. Areas containing immune-stained tissues were used for evaluation regardless the intensity of staining **(Simonsson et al., 2017)**.

### 4.9. Statistical analysis

Statistical analysis for numerical data was done using SPSS (IBM corp., Version 20). The mean number and the standard deviation (SD) were determined for parameters in each group. The significance of differences observed was assessed by ANOVA test followed by post HOC test. Significance was determined when probability (p-value) ≤ 0.05.

